## Supplementary information for "Encoding extracellular modification of artificial cell membranes using engineered self-translocating proteins"

General Information

Lipids:
1,2-dioleoyl-sn-glycero-3-phosphocholine (18:1 PC; DOPC) (Cat# 850375) was purchased from Avanti Polar Lipids in chloroform. Cholesterol (Cat# 700000P) was also purchased from Avanti Polar Lipids.

Protein expression and purification reagents:
PURExpress® kit was purchased from NEB (Cat# E6800), myTXTL® T7 Expression Kit from Daicel Arbor Biosciences (Cat# 505024). BL21(DE3) cells were purchased from NEB (Cat# C2527). HisPur™ Ni-NTA Resin was purchased from ThermoFisher (Cat# 88221). PD SpinTrap G-25 columns were purchased from Cytiva (Cat# 28918004).

Antibodies:
DyLight™ 650-Anti-6XHis-tag antibody was purchased from ThermoFisher (Cat# MA1-21315-D650).
CF™ 647-Anti-Somatostatin antibody was purchased from biorbyt (Cat# orb500667-CF647).
AF® 647-Anti-GLP1 antibody was purchased from Bioss (Cat# bsm-0933M-A647).

Ficoll® 400 was purchased from Millipore Sigma (Cat# F8016). Mineral oil heavy from Fisher Scientific (O122-1) and BSA (Bovine serum albumin) was purchased from Cytiva (SH30574.01). Glucose Oxidase (Cat# G2133) as well as catalase (Cat# C9322) were purchased from Millipore Sigma. Confocal microscopy was done on a Zeiss Cell Observer® SD consisting of a Yokagawa spinning disk system (Yokagawa, Japan) built around an Axio Observer Z1 (Zeiss, Germany) motorized inverted microscope. Microscopy images were analyzed in Fiji/ImageJ. Fluorescence values of individual GUVs are mean fluorescence values generated by manually selecting GUVs in ImageJ with the oval selection tool. For quantification, at least 10 GUVs from each population were averaged.

### Supplementary Fig. 1: Negative controls for antibody assay.

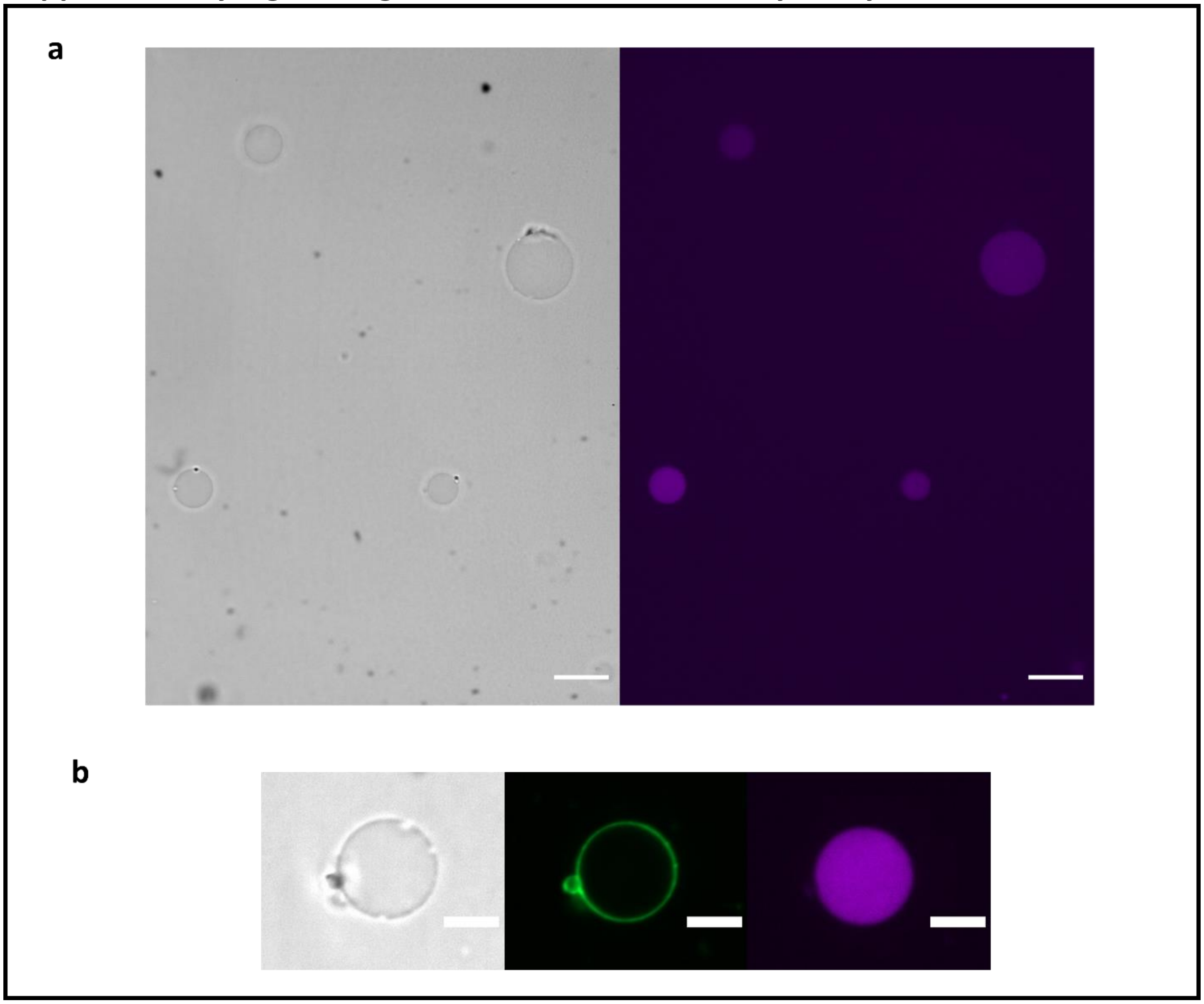


**a**, GUVs encapsulating Cy5 anti-6XHis-tag antibody show no membrane localization of antibody without αHL treatment after 2 hours. Scale bar: 25 µm **b**, GUVs encapsulating Cy5 anti-6XHis-tag antibody show no membrane localization of Cy5-antibody after 4 h treatment with αHL with $L_2$-GLP1-$L_2$ insert. Scale bar: 10 µm

**Supplementary Fig. 2: Cryo-EM 2D class averages of αHL with $L_2$-GLP1-$L_2$ insert.**

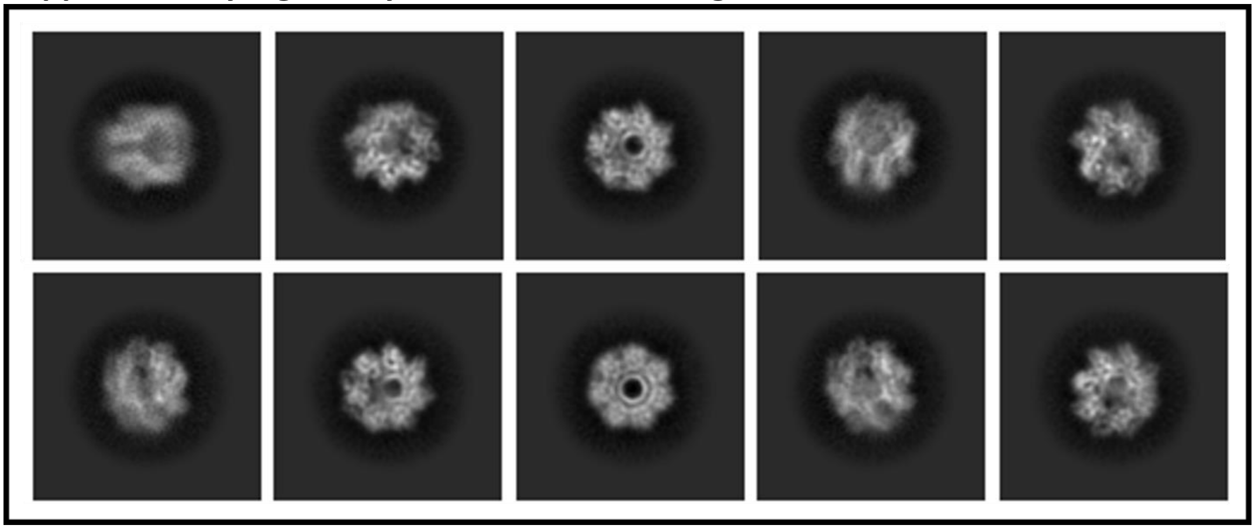

αHL monomer with the $L_2$-GLP1-$L_2$ insert was treated with sodium deoxycholate (6.25 mM) to induce pore formation.[48,49] The pore solution was diluted to 1 mM sodium deoxycholate and used for the preparation of Cryo-EM grids. 2D class-averages were generated from 36,740 particles collected from 5357 exposures. These 2D class-averages further confirm that peptide inserts in the loop region do not significantly alter the 3D structure of the αHL pore. Due to the high flexibility of the GGGGSGGGGS linkers, it is not possible to visualize the insert in the loop through cryo-EM.

### Supplementary Fig. 3: Leakage assay for αHL with K3 and E3 insert.

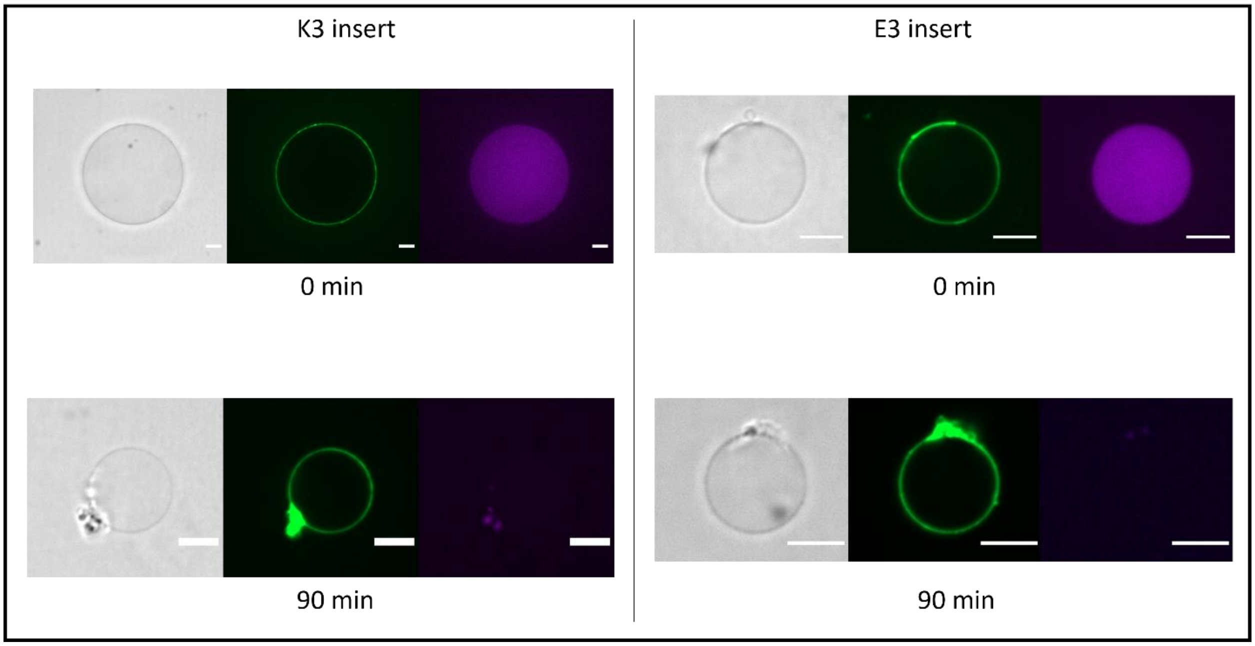


GUVs encapsulating Cy5 were formed through the inverse emulsion method with a lipid composition of 60% DOPC and 40% cholesterol. Addition of a GFP- αHL fusion protein with the K3 insert or E3 insert leads to leakage of Cy5 from the GUVs over time, showing that αHL with the K3 insert or E3 can assemble into functional pores. Shown are representative GUVs right after protein addition and after 90 minutes of incubation at RT. Leakage assays for αHL with K3 and E3 inserts were performed under low salt conditions. Scale bar: 10 µm

**Supplementary Fig. 4: Sufficient protein expression is necessary for tissue formation.**

a

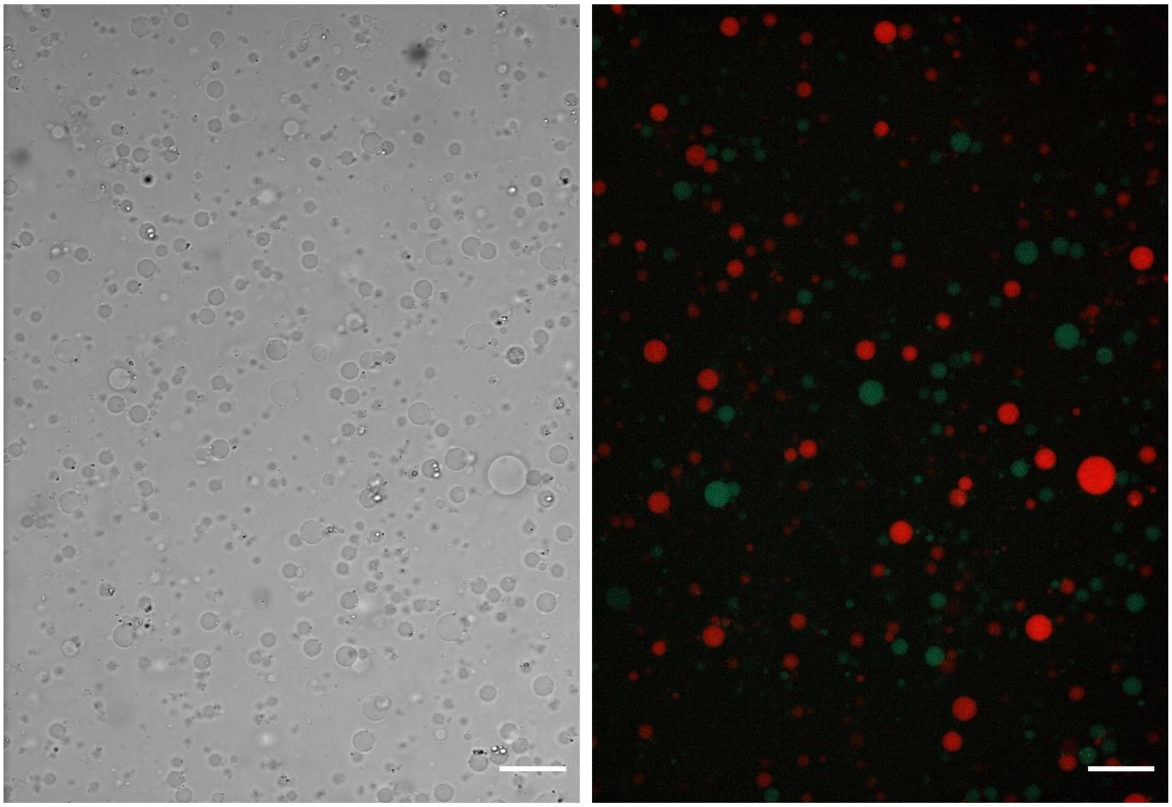

**a**, GUVs expressing αHL with either the K3 or the E3 insert were formed through the inverse emulsion method with a lipid composition of 60% DOPC and 40% cholesterol. The GUVs encapsulate the PURExpress system with a plasmid for expression of the αHL protein. For GUVs expressing αHL with the E3 insert, CFP was added to enable fluorescence imaging while for GUVs expressing αHL with the K3 insert mCherry was co-encapsulated. After incubation for 20 minutes at 37 °C no aggregation could be observed. Aggregation requires protein expression for at least one hour. Scale bar: 25 µm

### Supplementary Fig. 5: GUVs expressing αHL with K3 insert do not self-aggregate.

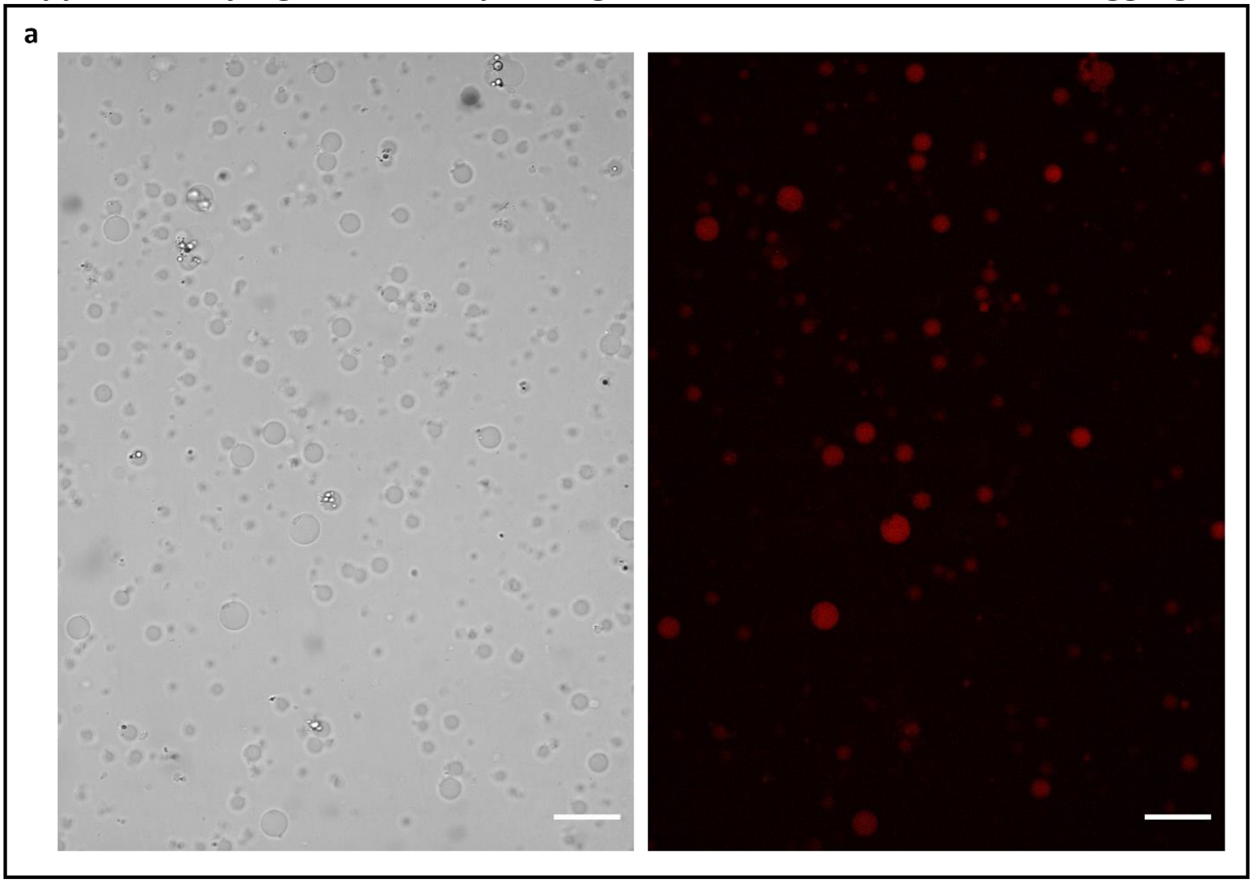

**a**, GUVs expressing αHL with the K3 insert were formed through the inverse emulsion method with a lipid composition of 60% DOPC and 40% cholesterol. The GUVs encapsulate the PURExpress system with a plasmid for expression of the αHL protein. Pre-expressed mCherry was added to enable fluorescence imaging. After protein expression for 2 hours at 37 °C no aggregation could be observed in the absence of GUVs expressing αHL with the E3 insert. Scale bar: 25 µm

### Supplementary Fig. 6: GUVs expressing αHL with E3 insert do not self-aggregate.

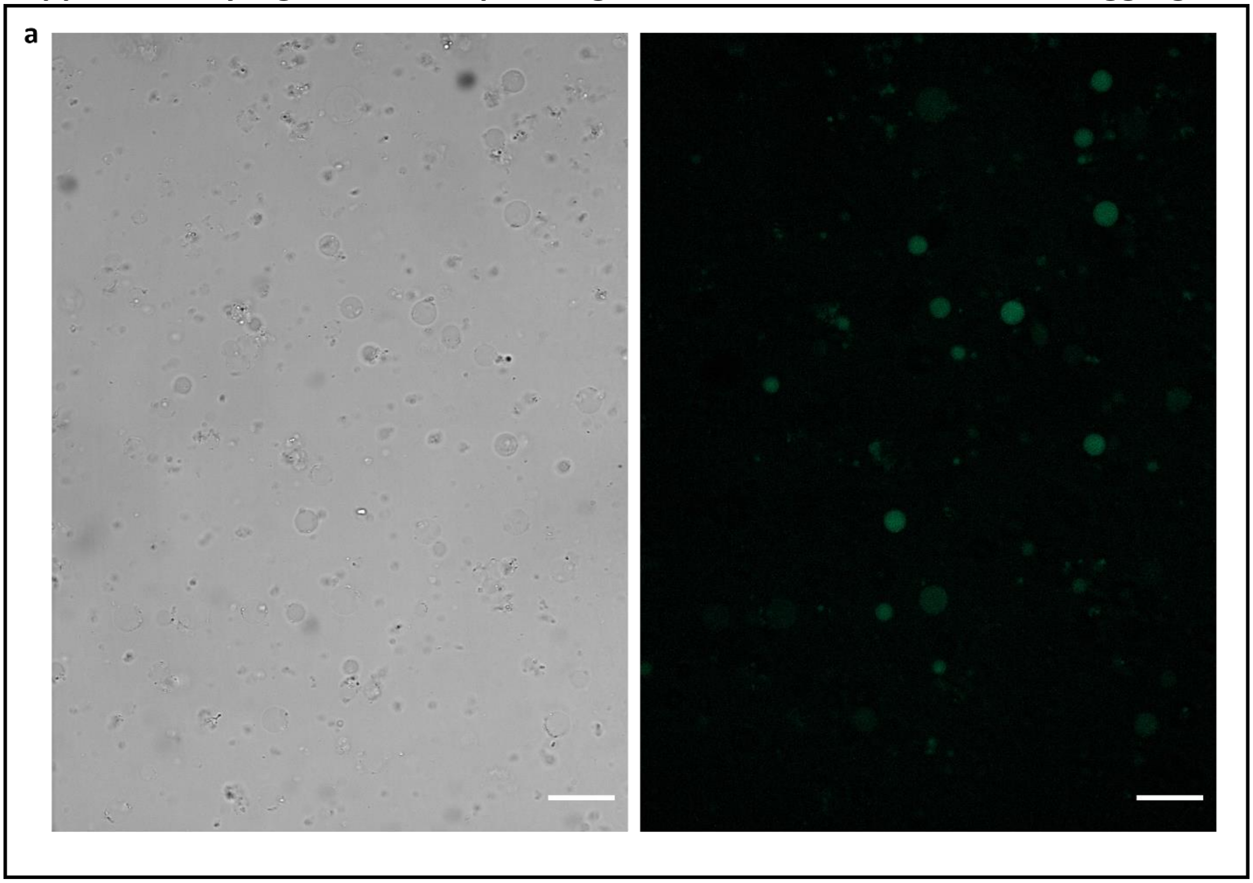


**a**, GUVs expressing αHL with the E3 insert were formed through the inverse emulsion method with a lipid composition of 60% DOPC and 40% cholesterol. The GUVs encapsulate the PURExpress system with a plasmid for expression of the αHL protein. Pre-expressed CFP was added to enable fluorescence imaging. After protein expression for 2 hours at 37 °C no aggregation could be observed in the absence of GUVs expressing αHL with the K3 insert. Scale bar: 25 µm

**Supplementary Fig. 7: Hyper7 fluorescence in tissue-like structures in the absence of glucose.**

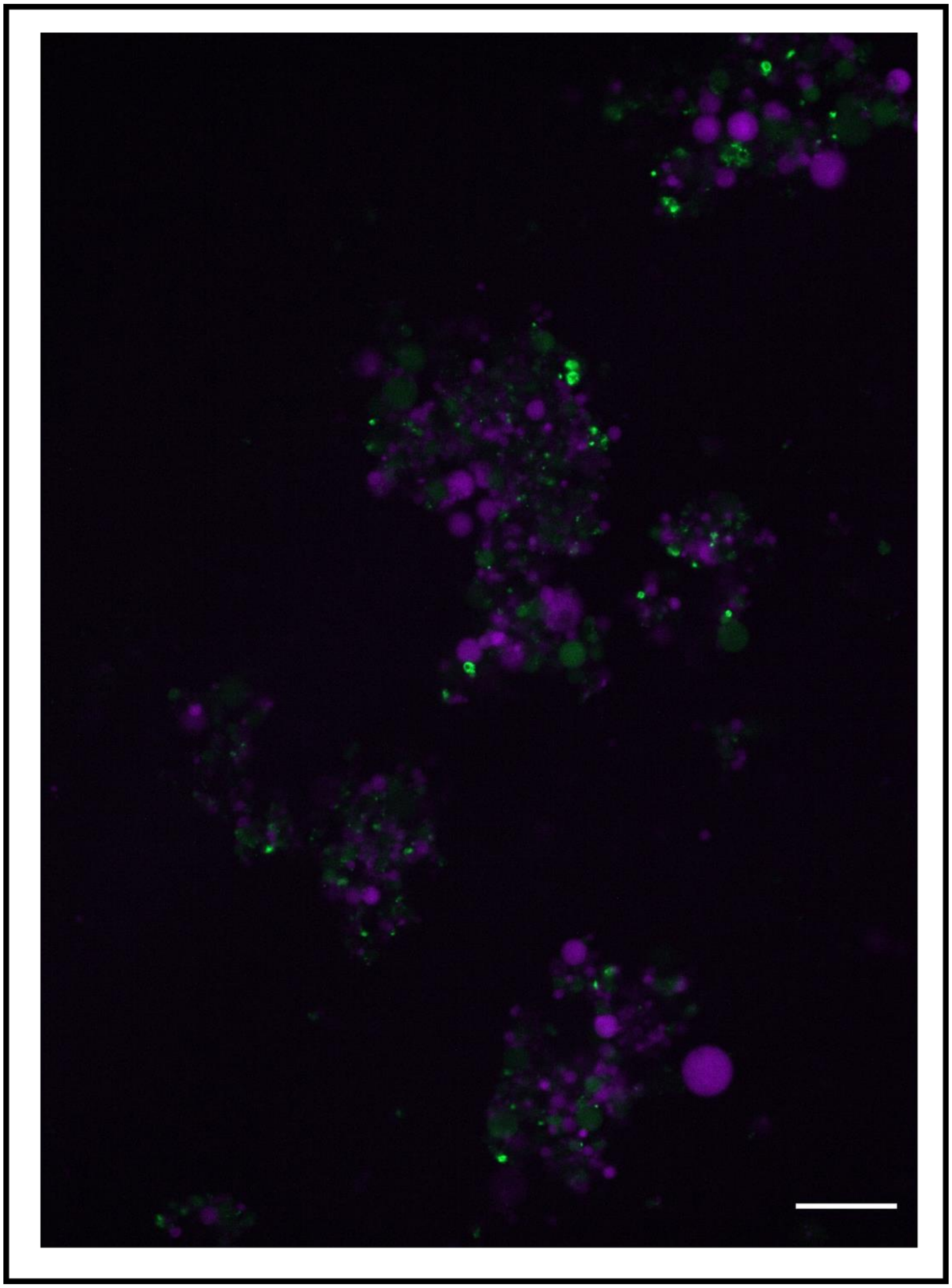

GUVs expressing αHL with either the K3 or the E3 insert were formed through the inverse emulsion method with a lipid composition of 60% DOPC and 40% cholesterol. For GUVs expressing αHL with the K3 insert, pre-expressed mCherry and glucose oxidase was co-encapsulated. For GUVs expressing αHL with the E3 insert, pre-expressed Hyper7 was co-encapsulated. The two populations of GUVs were mixed and incubated for 2 hours at 37 °C to enable expression of the respective αHL proteins. αHL expression led to the formation of tissue-like structures. In this control experiment, no glucose was added meaning that while glucose oxidase was present in the resulting tissue like structures, no significant formation of hydrogen peroxide was to be expected. As a result, Hyper7 fluorescence in these tissue-like structures is lower than in tissue-like structures where synthesis of hydrogen peroxide was induced by addition of glucose (Fig. 6).

**Supplementary Table 1: Peptide sequences which could be inserted into the αHL loop without disrupting its activity.**

| Insert | Amino Acid Sequence | Length |
|---|---|---|
| **L-6XHis-L** | GGGGS HHHHHH GGGGS | 16 |
| **$L_2$-6XHis-$L_2$** | $(GGGGS)_2$ HHHHHH $(GGGGS)_2$ | 26 |
| **$L_3$-6XHis-$L_3$** | $(GGGGS)_3$ HHHHHH $(GGGGS)_3$ | 36 |
| **L-Somatostatin-L** | GGGGS AGCKNFFWKTFTSC GGGGS | 24 |
| **$L_2$-GLP1-$L_2$** | $C(GGGGS)_2$ HAEGTFTSDVSSYLEGQAAKEFIAWLVKGR $(GGGGS)_2C$ | 52 |

Inserts listed in Supplementary Table 1 were cloned into the membrane translocating loop of αHL between D128 and K131, with the inserted peptide replacing T129 and G130. αHL proteins with these inserts proved to be fully functional, being able to self-insert into lipid bilayers and assembling into pores while translocating the respective insert across the membrane.

**Supplementary Table 2: Peptide inserts which lead to non-functional αHL proteins.**

| Insert | Amino Acid Sequence | Length |
|---|---|---|
| **L-Sac7e-L** | GGGGS<br>MAKVRFKYKGEEKEVDTSKIKKVWRVGKMVS<br>FTYDDNGKTGRGAVSEKDAPKELMDMLARA<br>EKKK GGGGS | 75 |
| **L-SNAP-L** | GGGGS<br>MDKDCEMKRTTLDSPLGKLELSGCEQGLHRIIF<br>LGKGTSAADAVEVPAPAAVLGGPEPLMQATA<br>WLNAYFHQPEAIEEFPVPALHHPVFQQESFTR<br>QVLWKLLKVVKFGEVISYSHLAALAGNPAATA<br>AVKTALSGNPVPILIPCHRVVQGDLDVGGYEG<br>GLAVKEWLLAHEGHRLGKPGL GGGGGS | 192 |
| **L-GFP-L** | GGGGS<br>MRKGEELFTGVVPILVELDGDVNGHKFSVRGE<br>GEGDATNGKLTLKFICTTGKLPVPWPTLVTTLT<br>YGVQCFARYPDHMKQHDFFKSAMPEGYVQE<br>RTISFKDDGTYKTRAEVKFEGDTLVNRIELKGID<br>FKEDGNILGHKLEYNFNSHNVYITADKQKNGI<br>KANFKIRHNVEDGSVQLADHYQQNTPIGDGP<br>VLLPDNHYLSTQSVLSKDPNEKRDHMVLLEFV<br>TAAGITHGMDELYK<br>GGGGS | 248 |

Inserts listed in Supplementary Table 2 were cloned into the membrane translocating loop of αHL between D128 and K131, with the inserted peptide replacing T129 and G130. αHL proteins with these inserts were non-functional, meaning that they did not form pores or insert into lipid-bilayers.
